# Phylogeography of equine infectious anemia virus

**DOI:** 10.1101/867747

**Authors:** Manuel Jara, Alba Frias-De-Diego, Gustavo Machado

## Abstract

Equine Infectious Anemia virus (EIAV) is the causative agent of one of the most impacting infectious diseases affecting equids. EIAV is genetically diverse with several lineages circulating globally. To elucidate EIAV global spread patterns, we studied its spatiotemporal dynamics applying Bayesian phylodynamic analyses, using a worldwide compiled dataset composed of unique sequences of the *gag* gene. In addition, we performed a scoping review of 1.470 publications on EIAV to characterize the spatiotemporal trends in EAIV research. Phylogeographic reconstruction suggested Hungary as the most likely country of origin for current EIAV circulation (root state posterior probability = 0.21), and one of the most important centers of diversification for the disease. Historical EIAV spread was predominantly characterized by long-distance spread across continents. As a result, we found that the American and Asian circulating EIAV (i.e., Mongolia) are more related to European lineages than to other Asian countries, being Europe the continent with the highest EIAV phylogenetic diversity. Our bibliometric analysis showed a continuous increment in the number of publications per year, where the United States and China appeared as the countries with highest EIAV-related scientific production. This study provides a historical geographic mapping of the EIAV linages’ spread patterns. In addition, we identified important asymmetry between the current research effort and the availability of genetic data which restricted our abilities to quantify for example the time frame of virus dispersal.

## INTRODUCTION

Equine infectious anemia (EIA) is a viral disease affecting equines that entails great economic losses worldwide mostly due to the obligatory elimination or isolation of infected animals (Montelaro et al., 1993; Souza, 2009; USDA, 2017). EIAV is currently one of the most significant infectious diseases of equids (R. F. Cook et al., 2013). It is caused by a lentivirus (EIAV) from the *Retroviridae* family (Leroux et al., 2004) known to be mostly transmitted by vectors from the family Tabanidae (Rojas, 1981; Barros and Foil, 2007). Due to the environmental needs of these vectors, EIA is more prevalent in warm and wet landscapes (Issel and Foil, 1984; Silva et al., 1999; Barros and Foil, 2007; Parr et al., 2014), but owing to a globalized animal trade and fast global warming, it has spread extensively through the globe in the last decades (Lima and Cintra, 2015). In the United States alone, the economic impact of EIA has been estimated to be US $38 million, which may be much bigger in countries with higher horse-related industry, such as Brazil (Diniz et al., 2017), known to have the largest population of equids in Latin America (Oliveira et al., 2017).

The field of evolutionary epidemiology has seen the implementation of integrative analyses of enormous multivariate datasets that encompass not only genetic and phenotypic data, but also geographic and ecological information. Besides, within the field of evolutionary epidemiology, molecular phylogenetics is increasingly becoming an accurate and impacting approach owing to its combination of genetic and environmental data to predict and control infectious diseases (Galvani, 2003; Mideo et al., 2008; Lion and Gandon, 2016). As a result, an increasing number of studies have finely quantified the factors and trajectories of lineage evolution over space and time. For example, Chu et al., (2013) revealed the spatial and phylogenetic radiation routes followed by Dengue virus worldwide. Later, Alkhamis et al., (2018) showed the phylodynamic spread patterns of African Swine Fever in Africa and Eurasia, while Jamal and Belsham, (2018), Omondi et al., (2019) and Jara et al., (2019), performed similar studies focused on the molecular evolution and epidemiology of Foot-and-Mouth Disease virus. The addition of spatial analyses to these methods has proven to be a remarkably useful tool to identify hot areas where treatment and control activities should be deployed. This combined approach has been widely applied to explain virulence, spread, invasion and species establishment of pathogens worldwide (Magiorkinis et al., 2009; Auguste et al., 2015; Pigeault et al., 2015; Weaver and Forrester, 2015; Dellicour et al., 2016; Rogalski et al., 2017; S Dellicour et al., 2017; Forni et al., 2018; Fountain-Jones et al., 2018). However, these methodologies are still not commonly applied for food animal diseases and therefore, major evolution and global diversification of several infectious disease remain an important bottleneck for better and universally applicable molecular testing (Dorey-Robinson et al., 2019), especially in diseases affecting horse populations, such as EIAV (Antolin, 2008; Struchiner et al., 2008; Dorey-Robinson et al., 2019).

Phylogenetic relationships of EIAV have been previously assessed by the analysis of outbreak cases reported worldwide (Nagarajan and Simard, 2007; Cappelli et al., 2011; Capomaccio et al., 2012), mainly focused in the areas with high horse-related industry, such as Brazil (Bicout et al., 2006; Nogueira et al., 2017; Barzoni et al., 2018; Cursino et al., 2018). However, the evolutionary and spread history of EIAV is still unknown (Dorey-Robinson et al., 2019).

In this study, we investigated the spatiotemporal dynamics of EIAV by using Bayesian phylodynamic analyses of a comprehensive genetic, geographical and temporal dataset regarding EIAV occurrences. In addition, we evaluated the global publication trends on EIAV, based on the assumption that temporal changes in research productivity may identify research priority needs. Results may be used to advance EAIV surveillance and support regional disease control policies related to intervention.

## METHODS

### Data collection and curation

We built a comprehensive genetic database by gathering information of the EIAV *gag* gene sequence (∼1,400 bp) along with the collection date and location as metadata (GenBank ID numbers and metadata available in Supplementary material). This gene was chosen due to its extended use in literature for phylogenetic studies (Olsen, 1998; Cappelli et al., 2011; Cook et al., 2013). Samples lacking any of the metadata information were discarded for subsequent analyses. This dataset comprises 297 sequences of 17 countries from 1947 to 2015. To determine phylogenetic relationship accurately, and to avoid sampling bias among EIAV sequences, we removed all duplicated sequences that represents the same outbreak. Thus, the cleaned database comprises 51 sequences.

### Discrete phylogeographical analysis

Sequences were aligned using Mega X, available at www.megasoftware.net (Kumar et al., 2008). The recombination detection program (RDP) v5.3 (Martin et al., 2015) was used to search for evidence of recombination within our dataset using five different methods (BootScan, Chimaera, MaxChi, RDP, and SiScan). Thus, four recombinant sequences were observed in EIA dataset.

To determine whether there was a sufficient temporal molecular evolutionary signal of the EIAV sequences used for each serotype phylogeny, we used TempEst v1.5 (Rambaut et al., 2016). To calculate the *P*-values associated with the phylogenetic signal analysis, we used the approach described by (Murray et al., 2016) based on 1,000 random permutations of the sequence sampling dates (Navascués et al., 2010). The relationship found between root-to-tip divergence and sampling dates (years) supported the use of molecular clock analysis in this study. The obtained results supported a significant temporal signal (*P*-value<0.05).

Phylogeographic history of EIAV dispersal was recovered from the obtained spatiotemporal phylogeny. The phylogenetic tree was generated by a discrete phylogeography estimation by Bayesian inference through Markov Chain Monte Carlo (MCMC), implemented in BEAST v2.5.0 (Bouckaert et al., 2014), applying a Hasegawa-Kishino-Yano (HKY+G; (Hasegawa et al., 1985)) substitution model with gamma-distributed rate heterogeneity among sites (Shapiro et al., 2006), to determine the best fitting model of evolution we used PhyML with Smart Model Selection (SMS) (Lefort et al., 2017). By using Nested Sampling Beast package v1.0.4 (Russel et al., 2019) we compared different molecular clock models to find the one that showed the best fit for our data. The marginal likelihood value supported the use of uncorrelated lognormal relaxed molecular clock (Drummond et al., 2006) using a constant population size model. To infer the epidemic demographic histories of EIAV we estimated the effective number of infections through time by using the Bayesian skyline plot approach (Drummond et al., 2005). All analyses were developed for 200 million generations, sampling every 10,000^th^ generation and removing 10% as chain burn-in. The Markov Chain Monte Carlo analysis was investigated using Tracer software v1.7 (Rambaut et al., 2018) to ensure adequate effective sample sizes (ESS) (above 200) which were obtained for all parameters. Final trees were summarized and visualized via Tree Annotator v. 2.3.0 and FigTree 1.4.3 respectively (included in BEAST v2.5.0) (Rambaut and Drummond, 2016; Rambaut, 2017).

To reconstruct the ancestral-state phylogeographic transmission across countries, we used the discrete-trait extension implemented in BEASTv2.5.0 (Rambaut and Drummond, 2016; Rambaut, 2017). In addition, to explore the most important historical dispersal routes for the spread of EIAV across countries, we used a Bayesian stochastic search variable selection (BSSVS) (Lemey et al., 2009). Using BSSVS approach, we identified and eliminated the nonzero rates of change between each pair of discrete traits (countries) based on its Bayes factor value obtained (lower than 3). To perform this analysis, a symmetric rate matrix was assumed (Drummond and Bouckaert, 2015). To infer the intensity of directional transitions (forward and backward) within a matrix of the discrete trait (countries), we used a Markov jumps approach. To interpret the Bayes factors, a value of <3 is not considered significant, BF= 3.1-20 represents positive support, BF= 20.1-150 represents strong support, while >150.1 represents an overwhelming support (Drummond and Bouckaert, 2015). Finally, to visualize the spatiotemporal diffusion of EIAV we used Spatial Phylogenetic Reconstruction of Evolutionary Dynamics using Data-Driven Documents (D3) SPREAD3 software (Bielejec et al., 2016).

### Global EIAV research trends

We developed an overview of the research done on EIAV through a scoping review by searching for publications with “equine infectious anemia” in the title using Scopus database. We captured information related about authors, author affiliation, journal, year, title, abstract and keywords. Bibliometric analysis was performed in R using the Bibliometrix package v2.2.0 (Aria and Cuccurullo, 2017). We assigned the publications to each country by searching for the country name within the corresponding author’s affiliation, title, and abstract. Publications with multiple countries listed within the title and abstract were assigned to all countries involved. All country name variants were standardized. Using this information, we mapped the scientific productivity per country. To visualize the average annual trends, we fitted a locally weighted regression (‘LOESS’) line (Cleveland, 1979). In addition, we visualized the multinational collaboration network of the 20 most productive countries on EIAV research. Finally, we extracted the total horse population per country (Gilbert et al., 2018) which was used to correlate with scientific production.

## RESULTS

We found a total of 297 EIAV *gag* gene sequences from the 17 countries. After removing all highly recombinant sequences, as well as the similar sequences that represent the same outbreak event, 47 sequences were used for the phylogeographical analysis (Figure 1, Supplementary Table S1 for more details).

**FIGURE 1.**
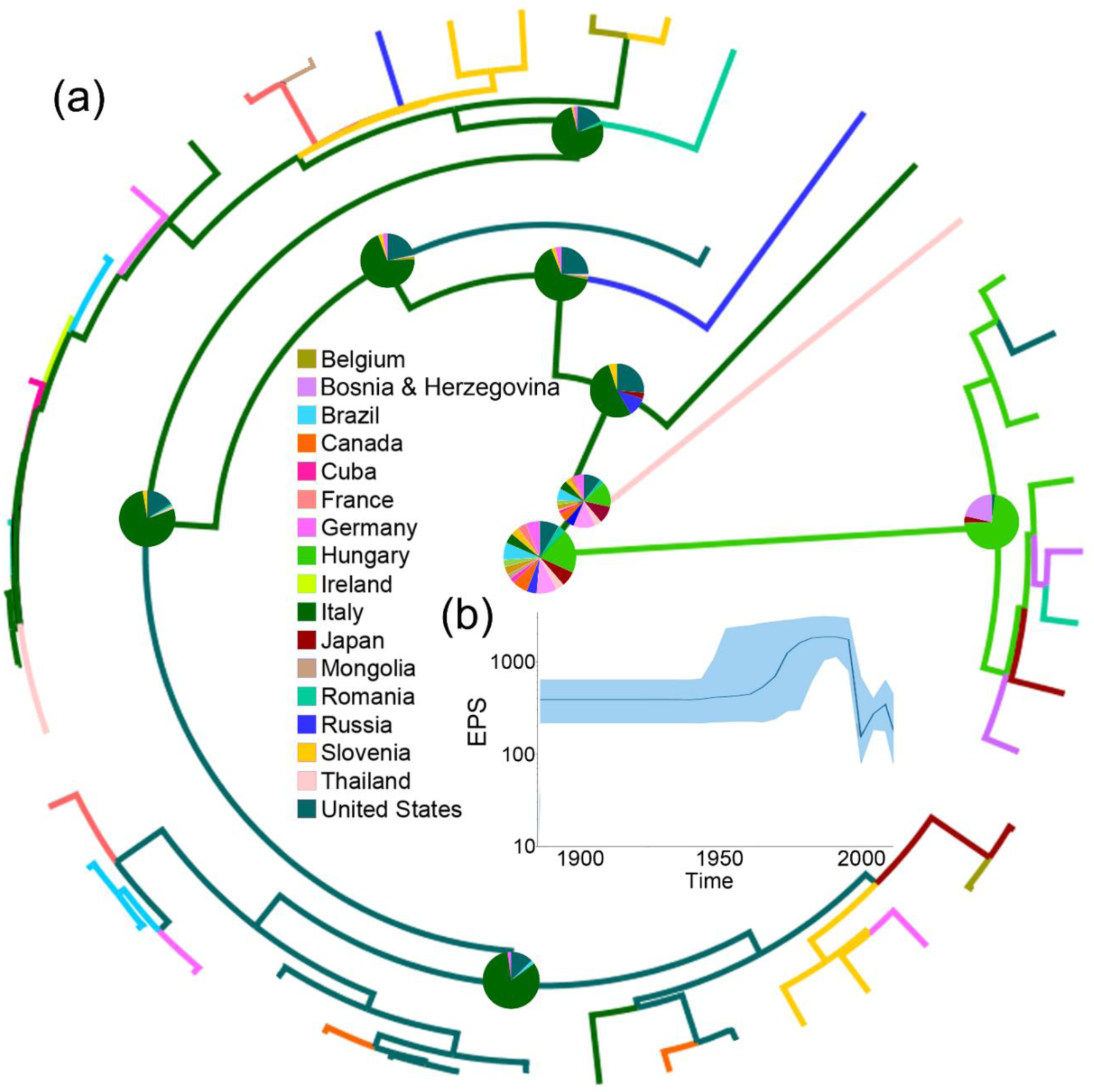
Dispersal history of EIAV lineages, as inferred by discrete phylogeographic analysis. (a) Maximum clade credibility phylogeny of EIAV colored according to the countries where the disease has been registered. The probabilities of ancestral areas (inferred from the Bayesian discrete trait analysis) are shown in pie charts. (b) Bayesian Coalescent Skyline analysis of EIAV epidemiological spread, here the darker line represents the median estimate of the estimated effective population size (EPS). The blue area shows the upper and the lower estimates of 95% interval.

### Phylogeography of equine infectious anemia virus

The phylogeographic reconstruction of EIAV indicated Hungary as the most likely origin of EIAV (root state posterior probability [RSPP] = 0.21). On the other hand, Italy and the United States represented the highest source of dispersal events, here represent regions from where a considerable number of lineages spread to other areas (Figure 1a). Bayesian skyline plot (BSP) showed a steady pattern in the genetic diversity (population size) until around 1970, when there was a sharp increase in its effective population size, evidencing a constant decrease around mid-1990s follow another increase in diversity in early 2000 (Figure 1b).

From early times on its evolutionary history, EIAV transmission has been characterized by its spread to distant regions (i.e. to countries not sharing international borders with the country of origin). This long-distance spread occurred in all directions, as seen in the case of Hungary, where the disease spread through Europe and Asia, including very distant countries, such as Russia, Japan, and Thailand. Likewise, we observed the outbreak occurred in United States in 1947, this event gave origin to the spread of EIAV across most of the American countries. On the other hand, Italy was the most likely responsible for the first introduction of the virus into Romania and Cuba (Figure 2a). Another important aspect of the spatiotemporal dynamics of EIAV was the multi-directionality of its spread, expressed as the movement of the disease in both directions between Hungary-United States and United States-Canada (see detailed footage of the transmission routes of EIAV in Supplementary video 1). Bayesian stochastic search variable selection (BSSVS-BF) results showed the most significant viral transmission routes for EIAV, where the most intense were represented from Germany to Japan (BF=16,080.9), followed by the spread from Belgium to the United States (BF=804.5), from Mongolia to Cuba (BF=402.2) and from Hungary to Ireland (BF=201.1) (Figure 3b).

**FIGURE 2.**
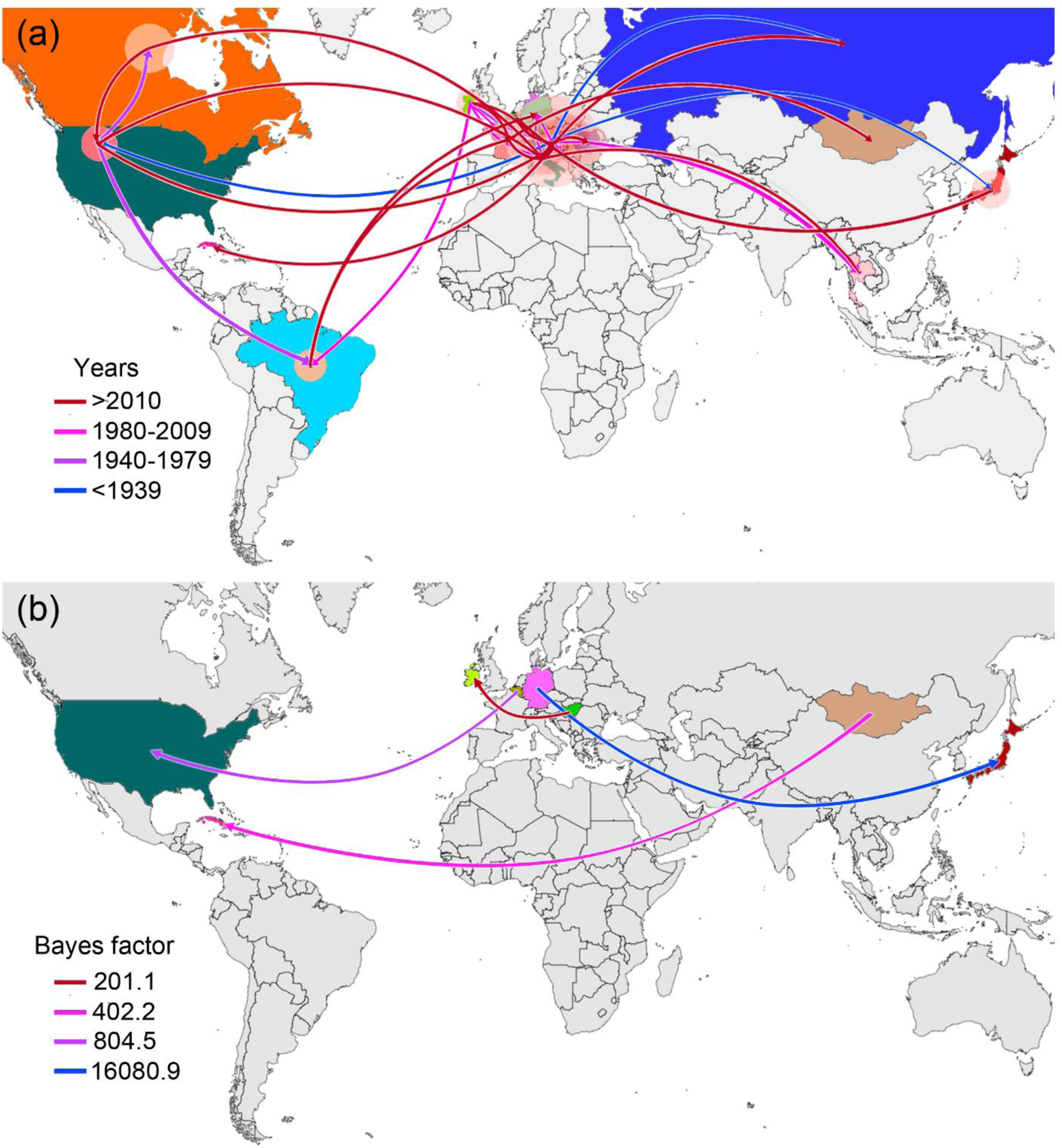
Global spatiotemporal diffusion of EIAV epidemic. (a) Phylogeographic patterns of EIAV spread where diameters of the colored circles are proportional to the number of maximum clade credibility (MCC) branches, maintaining a particular location state at each time period. The color of the branches represents the age of the internal nodes, where darker red colors represent older spread events. (b) Representation of the most significant location transitions for EIAV spread based on only the rates supported by a BF greater than 3 are indicated, where the color of the branches represents the relative strength by which the rates are supported.

**FIGURE 3.**
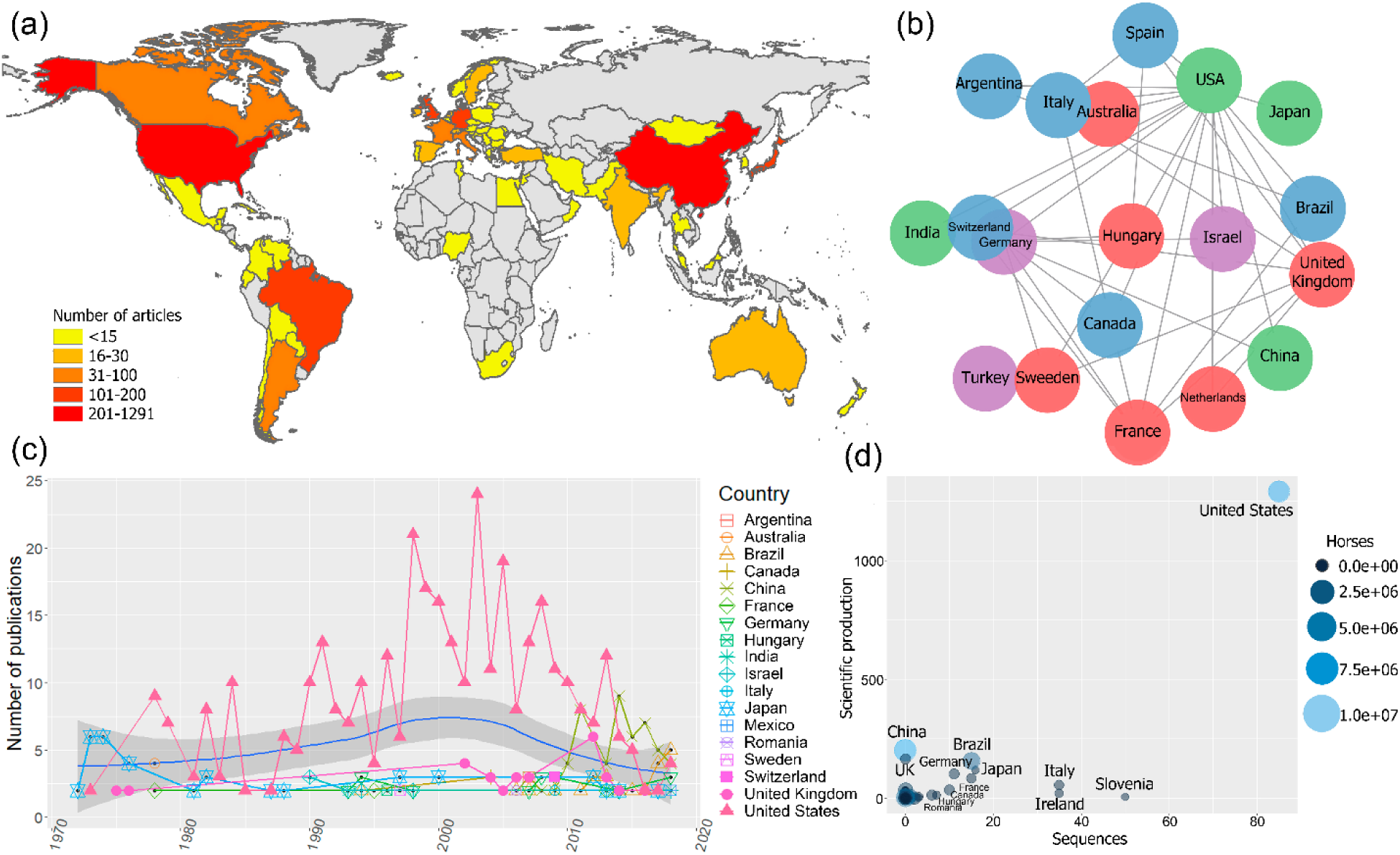
Global scientific production on EIAV. (a) Scientific production on EIAV measured by each country. (b) Multinational collaboration network based on the 20 most productive countries, where clusters of similar colors indicate the countries with a higher tendency to collaborate with one another in scientific publications. (c) Total number of publications per year, blue solid line indicates the average long-term trends in the scientific productivity (annual LOESS fits). (d) Bubble chart showing the relationship between the number of sequences, scientific production and the total number of horses per country (source of horse population (Gilbert et al., 2018).

### Scientific production on equine infectious anemia virus

Bibliometric analyses resulted in 1.470 unique publications from 1946 to 2019, from 57 countries. There were countries with a minimum of 1 and maximum of 1.291 publications, the most productive countries were the United States, China and the United Kingdom (Figure 3a).

The global collaboration network resulted from the 20 most productive countries, revealed that the United States had the largest collaboration network, followed by the United Kingdom, Germany, and France. Asian countries and Australia had the lowest levels of multinational research within this group, and we did not observe any African countries on this list, which indicates their low levels of international collaboration compared to the rest of continents (Figure 3b). Bibliometric analysis also showed that the research effort over time appeared to be relatively constant from 1946 until mid-1960. From this point on there is a clear increment in the scientific productivity which also vary substantially over the time. However, during the period 1980-1985 we observed a reduction in the number of published works. Overall, the average annual growth rate of publications was 23.3%, and the most productive year was 2013 with 59 articles (Figure 3c). The average annual global trends in EIAV scientific productivity showed a Gaussian shape in the number of publications, with a peak of productivity on 2000, after that, we observe a decreasing tendency that still lasts until today (Figure 3c). We also observed that around mid-1970s, Japan was the country with more publications on EIAV. Later, from 1978, United States started consolidating its position as the country with the highest number of publications in the majority of years, showing its highest research effort around 2002. However, from 2007, the scientific productivity of United States showed a continuous decrease, and was surpassed in 2013 by China, that obtained the first position, and later in 2018, Brazil appeared to be on top of the global research productivity on EIAV (Figure 3c).

Unsurprisingly, the highest scientific productivity was related to the countries with largest horse population (i.e., China, United States, and Brazil), being United States the country that exhibited the highest values in almost all the categories. United States showed the highest scientific production on EIAV (Figure 3a), accompanied by the highest number of EIAV genetic sequences, as well as and the most complex collaboration network (Figure 3a-d). China, which is the country with the largest horse population and the second most productive country in terms of scientific articles on EIAV but had not yet contributed to the pool of available sequences (as needed for phylogeographic analysis, see methods for detailed information). However, countries like Hungary, Italy and Ireland, which have a small horse population and little scientific productivity, showed a big amount of publicly available sequences which contributed to the phylogeographic analysis (Figure 3d).

## DISCUSSION

This study represents an attempt to investigate the global spatiotemporal patterns of EIAV from a multidisciplinary perspective. We combined spatial and evolutionary approaches with a scoping review framework to provide an overview of the historical knowledge of the disease. We reconstructed the evolutionary history of the available lineages of EIAV along with the most likely origin of the analyzed sequences, the most relevant countries that have been recognized as important centers of virus diversification, and the most important spread routes. We quantified the global patterns of the scientific literature.

### Phylodynamic patterns of equine infectious anemia virus spread

Equine Infectious Anemia virus is widely distributed throughout the world (Cook et al., 2001). Our phylogeographic results suggested Hungary was most likely center of origin for the currently available sequences. From this country, EIAV diversified in several directions across the globe. The most similar lineages with the ancestral strains come from Italy, supporting what was previously described by Kuhar and Malovrh (2016). Furthermore, as previously discover by Capomaccio et al., (2012a,b), there was an important spread route of EIAV from Europe to the Americas, which could be directly related to the intensive movement of animals between these continents. Likewise, Tigre et al., (2017), found high similarities among Brazilian EIAV isolates and European and North American strains.

Similarly to the historical spread exhibited by other animal diseases, such as Foot and Mouth Disease (Bachanek-Bankowska et al., 2018) or *Avian avulavirus* (Hicks et al., 2019), EIAV has shown the ability to spread rapidly through long distances, with evident jumps across continents. Spatiotemporal patterns of EIAV visualized through skyline plot showed that its worldwide spread have been accompanied by a constant increase in the genetic diversity that started around 1970 and ceased 20 years later, evidencing an abrupt decrease that could be explained by the intensification of disease control programs in countries with elevated count cases and the development of better ELISA-based EIAV serologic tests (Issel and Cook, 1993). For example, in United States, the improvements in the surveillance methods have allowed an increase in the number diagnostic tests from 2,777 in 1972 to 993,712 in 1991 (Issel et al., 2013). However, during the last decade, we observed some irregular trends in the genetic diversity of the virus, as a result of numerous outbreaks occurred worldwide, causing dramatic economic consequences (Timoney, 2000; Cook et al., 2013; Herholz et al., 2013). These epidemic pattern has been facilitated by the continuously growing international horse trade (Metcalf, 2001, Dominguez et al., 2015).

Based on our results, it is possible to observe a high genetic diversity of the virus, especially in Europe, where the countries surrounding two of the major centers of diversification (Hungary and Italy) accumulated the highest viral diversity, which supports the findings previously described by Kuhar and Malovrh (2016), and Gaudaire et al. (2018). Our results showed that the spread of EIAV into the Americas occurred due to the arrival of a strain from the United States, which later diversified and spread through Canada and Brazil. These results support a previous study of Tigre et al., (2017) that evidenced strong similarities between the Brazilian strains with the ones circulating in the United States. Moreover, our results also showed that most of the American strains returned to Europe, increasing the similarities between the strains found in these areas. On the other hand, the colonization of Asia seemed to occur exclusively from the arrival of European lineages, which explains the previously reported high similarities between certain strains from both continents in comparison with other lineages from Asia or the Americas (Herholz et al., 2013; Sharav et al., 2017). Finally, while the genetic data used here comes from hotspots areas of EIAV, there are still important gaps of information exemplified in the absence of genetic information in areas such as Africa, where despite it has evidenced the presence of three endemic EIAV serotypes (Herholz et al., 2013; Alnaeem and Hemida, 2019), there is a remarkable lack of genetic information, which could be highly related to an underreporting or lack of laboratory capacities which should be advocated (Barzoni et al., 2018).

### Global trends in equine infectious anemia virus research

The bibliometric analysis synthesised the global research trends relevant to map areas that have produce more scientific publications on EIAV. We found that overall, in opposition to Lotka’s law (Coile, 1977), the average number of scientific publications per year showed a steady growth until 2000, followed by a slow reduction maintained until today. However, China and Brazil break this pattern by showing a remarkable increase in their scientific production since around 2005 (Fig. 4c). Our results also highlighted the global importance of United States and China as the countries with more publications on EIAV. These results were unsurprising since both of them are known to be within the top countries in scientific production worldwide (Zhou and Leydesdorff, 2006; Nafade et al., 2018). It is well known that variations in scientific productivity between countries can be related to differences in factors such as language, training, development, and funding among others (Ramos et al., 2004), which we propose is one of the main reasons influencing the ability of less-developed countries to carry out the necessary scientific research studies to detect and report EIAV. We observed some clear examples of this in countries like Mexico, Ethiopia, Russia, and Mongolia, which despite being in the top 10 of the countries with higher horse population, showed very low scientific production about EIAV. Likewise, as in previous bibliometric research (Vanni et al., 2014; Ekundayo and Okoh, 2018) we observed asymmetries in the multinational alliances between developing and developed countries that have been historically endemic for EIAV (i.e., African countries).

Overall, our results showed that historically, the United States has been the most productive country which is strongly related to the fact that is one of the countries with highest horse population worldwide, being only surpassed by China (FAO, 2019). Another important aspect to highlight is the complex collaboration network that the United States shows, which has been historically considered as an important factor for the country scientific success. As demonstrated by previous bibliometric analyses that have shown a continuous increment in the multinational scientific collaboration over the past two decades (Cronin et al., 2004; Cronin, 2005; Sonnenwald, 2007), along with a direct link between scientific collaboration and the impact of the produced research (Figg et al., 2006). These differences in collaboration and impact can be observed in a number of Asian countries (i.e., China, Japan, Turkey, and India), which interestingly were observed as some of the less connected countries to the multinational collaboration network and yet were seen to be part of the most productive in relation to EIAV, although the impact of their research tends to be lower.

During the last five years, the patterns on the publication rates of the different countries have not followed a similar pattern. We observed that most of the countries, including the United States, have shown a reduction in the number of publications on EIAV, which started on 2003. On the other hand, we observed that China and Brazil increased their scientific productivity with more publications on this topic worldwide, tendency that remained until 2018. This change may be due to the increase on biosafety awareness and the need to understand how to control a disease that directly affects countries’ economies, since they are among the countries with highest equine-related animal trade worldwide. Also shows that countries such as the United States lower their research priorities in the past few years.

### Limitations

A relevant limitation encountered during the development of this study was caused by the lack of genetic data and inconsistencies in data availability, which introduced bias to our phylodynamic results (Xia, 2014). This is evidence by the under-representations of some countries with known and long history of being endemic for EIAV form which either no genome sequences were found, or the amount of available information was limited. Similarly, countries such as China, South Africa or Nigeria have published information about the disease, but no genetic data (including date of sample collection) are available. These inconsistencies are one of the major factors impeding the development of more accurate analysis assessing EIAV and may influence the results obtained in our phylogeographic analyses, which might need to be updated as more genetic dataset become available. This lack of information is hindering the development of optimal control strategies to minimize the impact of EIAV and other infectious diseases worldwide. Furthermore, this study also has some limitations related to the bibliometric section. (i) We only used publications from Scopus database, and (ii) the countries analysed as the locations where research was conducted were based on the affiliation address of the first author, which does not necessarily reflect that information, especially in articles with multinational collaborations.

## Conclusions

This study illuminate the EIAV global evolution and the main disproportion between occurrence of EIAV and presence of genetic information. This study suggested that Hungary was likely the origin EIAV and also acted as main source for global dispersal. The most significant viral transmission routes were from Germany to Japan. Here we identified countries with great importance for EIAV spread, consequently our results can be used to redirect resource allocation such as those dedicated to expanding molecular diagnostic and disease surveillance. Importantly, we highlight the importance of the genetic characterization and detailed report of the virus in future outbreaks. In addition, the incorporation of a bibliometric analysis provided a holistic view of the historical global research efforts on EIAV, ultimately provided evidence of knowledge gaps which can be used for example by local stakeholder while defining disease control programs and prioritize funding for future research.

## ACKNOWLEDGEMENTS

We acknowledge the Department of Population Health and Pathobiology-North Carolina State University provided startup funds for G. Machado and M. Jara. We also acknowledge RISF and CVM-NCSU for funding this project partially.

## Supplementary material

**Table S1.** Sample information for all Equine Infectious Anemia virus sequences used in this study.

**TABLE S2.** Global relevant information of Equine infectious Anemia at country level.

**Figure S1.** Treemap plot showing the number of publications on EIAV per subject area.

