## supplementary information for "Phylogeography of equine infectious anemia virus"

**SUPPLEMENTARY MATERIAL**

**TABLE S1.** Sample information for all Equine Infectious Anemia virus sequences used in this study.

| GeneBank accession ID | Country | Collection year |
| --- | --- | --- |
| JX003262 | Belgium | 2011 |
| JX193070 | Belgium | 2010 |
| GQ996594 | BosniaAndHerzegovina | 2009 |
| GQ996595 | BosniaAndHerzegovina | 2001 |
| JX480681 | Brazil | 2006 |
| KC213779 | Brazil | 2011 |
| KC213789 | Brazil | 2012 |
| AY968061 | Canada | 2005 |
| EF418579 | Canada | 2007 |
| HM177437 | Cuba | 2010 |
| KM248275 | France | 2014 |
| KT764944 | France | 2009 |
| AB693824 | Germany | 2012 |
| KC213790 | Germany | 2011 |
| KF878274 | Germany | 2013 |
| GQ996589 | Hungary | 2008 |
| GQ996592 | Hungary | 2009 |
| GQ996598 | Hungary | 2001 |
| JX193072 | Ireland | 2010 |
| JX480668 | Ireland | 2006 |
| EF418585 | Italy | 2007 |
| GQ927489 | Italy | 2006 |
| GQ927504 | Italy | 2008 |
| HM177435 | Italy | 2010 |
| HQ853234 | Italy | 2009 |
| AB675093 | Japan | 2011 |
| AB693823 | Japan | 2013 |
| JQ627111 | Japan | 2009 |
| KT764956 | Mongolia | 2009 |
| GQ229581 | Romania | 2009 |
| GQ927497 | Romania | 2008 |
| KF977825 | Russia | 2012 |
| KM202106 | Russia | 2011 |
| KF878275 | Slovenia | 2013 |
| KF977802 | Slovenia | 2009 |
| KF977821 | Slovenia | 2011 |
| KF977824 | Slovenia | 2010 |
| EU741609 | Thailand | 2006 |
| FJ713789 | Thailand | 2008 |
| AB693825 | USA | 1947 |
| AF172133 | USA | 1999 |
| AF330039 | USA | 2000 |
| AY742253 | USA | 2004 |
| AY968059 | USA | 2005 |
| LC178575 | USA | 2015 |
| M87575 | USA | 1992 |

**TABLE S2.** Global horse production and scientific production on Equine infectious Anemia at country level.

| Country | Total number of horses | Number of publications on EIAV |
| --- | --- | --- |
| United States | 10510748 | 1291 |
| Slovenia | 22790 | 5 |
| Italy | 367561 | 56 |
| Ireland | 84900 | 21 |
| Japan | 14722 | 119 |
| France | 367162 | 84 |
| Brazil | 5501872 | 157 |
| Germany | 448146 | 103 |
| Canada | 398303 | 35 |
| Hungary | 53000 | 14 |
| Romania | 470331 | 14 |
| Belgium | 29531 | 8 |
| Bosnia and Herzegovina | 15599 | 0 |
| Thailand | 6500 | 6 |
| Russia | 1381331 | 0 |
| Cuba | 888800 | 5 |
| Mongolia | 3939813 | 4 |
| China | 11019575 | 201 |
| United Kingdom | 409288 | 166 |
| Argentina | 2527764 | 35 |
| Israel | 4000 | 34 |
| Switzerland | 55535 | 28 |
| Australia | 264627 | 27 |
| Turkey | 120040 | 26 |
| India | 625453 | 23 |
| Spain | 260961 | 20 |
| Sweden | 102515 | 20 |
| Netherlands | 140566 | 19 |
| Austria | 90354 | 12 |
| Croatia | 23721 | 12 |
| Colombia | 1185896 | 11 |
| South Korea | 28235 | 8 |
| Greece | 27604 | 6 |
| Portugal | 36963 | 6 |
| Venezuela | 522000 | 5 |
| Poland | 185494 | 4 |
| Czechia | 34548 | 3 |
| Mexico | 6380179 | 3 |
| Pakistan | 400000 | 3 |
| Bolivia | 502502 | 2 |
| Ecuador | 209990 | 2 |
| Iceland | 64679 | 2 |
| Iran | 132815 | 2 |
| Lithuania | 16324 | 2 |
| Norway | 33985 | 2 |
| Moldova | 36800 | 2 |
| Serbia | 16560 | 2 |
| South Africa | 322771 | 2 |
| Bulgaria | 42725 | 1 |
| Chile | 300000 | 1 |
| Costa Rica | 127922 | 1 |
| Egypt | 75730 | 1 |
| Gambia | 6000 | 1 |
| Guatemala | 132132 | 1 |
| Jordan | 2247 | 1 |
| Malaysia | 4217 | 1 |
| Nigeria | 102436 | 1 |
| Paraguay | 275000 | 1 |
| Tunisia | 57254 | 1 |
| Afghanistan | 175000 | 0 |
| Albania | 32258 | 0 |
| Algeria | 46841 | 0 |
| Angola | 1006 | 0 |
| Antigua and Barbuda | 491 | 0 |
| Armenia | 10631 | 0 |
| Azerbaijan | 68520 | 0 |
| Barbados | 1265 | 0 |
| Belarus | 49000 | 0 |
| Belize | 5978 | 0 |
| Benin | 701 | 0 |
| Bermuda | 1018 | 0 |
| Bhutan | 13322 | 0 |
| Botswana | 35210 | 0 |
| British Virgin Islands | 100 | 0 |
| Burkina Faso | 41163 | 0 |
| Cabo Verde | 560 | 0 |
| Cambodia | 30486 | 0 |
| Cameroon | 17823 | 0 |
| Chad | 1166908 | 0 |
| Congo | 79 | 0 |
| Cook Islands | 303 | 0 |
| Cyprus | 661 | 0 |
| North Korea | 48266 | 0 |
| Denmark | 51586 | 0 |
| Dominican Republic | 359016 | 0 |
| El Salvador | 98017 | 0 |
| Estonia | 6300 | 0 |
| Eswatini | 1689 | 0 |
| Ethiopia | 2228258 | 0 |
| Falkland Islands | 1147 | 0 |
| Fiji | 47429 | 0 |
| Finland | 74400 | 0 |
| French Guiana | 860 | 0 |
| French Polynesia | 2200 | 0 |
| Georgia | 39636 | 0 |
| Ghana | 2947 | 0 |
| Greenland | 140 | 0 |
| Grenada | 30 | 0 |
| Guadeloupe | 69 | 0 |
| Guam | 63 | 0 |
| Guinea | 3450 | 0 |
| Guinea-Bissau | 2467 | 0 |
| Guyana | 2418 | 0 |
| Haiti | 503489 | 0 |
| Honduras | 181244 | 0 |
| Indonesia | 442602 | 0 |
| Iraq | 50668 | 0 |
| Jamaica | 4008 | 0 |
| Kazakhstan | 2415654 | 0 |
| Kenya | 2095 | 0 |
| Kuwait | 1147 | 0 |
| Kyrgyzstan | 481329 | 0 |
| Laos | 32550 | 0 |
| Latvia | 9278 | 0 |
| Lebanon | 3261 | 0 |
| Lesotho | 64410 | 0 |
| Libya | 45476 | 0 |
| Liechtenstein | 263 | 0 |
| Luxembourg | 4725 | 0 |
| Madagascar | 498 | 0 |
| Malawi | 87 | 0 |
| Mali | 561500 | 0 |
| Malta | 1064 | 0 |
| Martinique | 968 | 0 |
| Mauritania | 66390 | 0 |
| Mauritius | 150 | 0 |
| Montenegro | 3784 | 0 |
| Morocco | 188000 | 0 |
| Myanmar | 99854 | 0 |
| Namibia | 45656 | 0 |
| New Caledonia | 11274 | 0 |
| New Zealand | 43684 | 0 |
| Nicaragua | 268399 | 0 |
| Niger | 250683 | 0 |
| North Macedonia | 17951 | 0 |
| Panama | 105465 | 0 |
| Papua New Guinea | 2044 | 0 |
| Peru | 749657 | 0 |
| Philippines | 248909 | 0 |
| Puerto Rico | 6123 | 0 |
| Qatar | 6074 | 0 |
| Reunion | 374 | 0 |
| Saint Helena | 0 | 0 |
| Saint Lucia | 1126 | 0 |
| Samoa | 1936 | 0 |
| Saudi Arabia | 28000 | 0 |
| Senegal | 550798 | 0 |
| Sierra Leone | 438794 | 0 |
| Slovakia | 6407 | 0 |
| Solomon Islands | 151 | 0 |
| Somalia | 885 | 0 |
| Sri Lanka | 1383 | 0 |
| Sudan | 790876 | 0 |
| Suriname | 297 | 0 |
| Syrian Arab Republic | 14050 | 0 |
| Tajikistan | 80408 | 0 |
| Timor Leste | 52916 | 0 |
| Togo | 1846 | 0 |
| Tonga | 11775 | 0 |
| Trinidad and Tobago | 1448 | 0 |
| Turkmenistan | 25840 | 0 |
| Ukraine | 291500 | 0 |
| United Arab Emirates | 438 | 0 |
| Uruguay | 420618 | 0 |
| Uzbekistan | 230600 | 0 |
| Vanuatu | 7630 | 0 |
| Viet Nam | 86759 | 0 |
| Yemen | 1857 | 0 |
| Zimbabwe | 28268 | 0 |
